# 1D confinement mimicking microvessel geometry controls pericyte shape and motility

**DOI:** 10.1101/2023.12.20.572195

**Authors:** Aude Sagnimorte, Marie R. Adler, Gaspard de Tournemire, Pablo J. Sáez, David Gonzalez-Rodriguez, Claire A. Dessalles, Avin Babataheri

**Affiliations:** LadHyX, CNRS, Ecole polytechnique, Institut polytechnique de Paris, 91120, Palaiseau, France; Cell Communication and Migration Laboratory, Institute of Biochemistry and Molecular Cell Biology, Center for Experimental Medicine, University Medical Center Hamburg-Eppendorf, Hamburg, Germany; Université de Lorraine, LCP-A2MC, F-57000, Metz, France; Department of Biochemistry, University of Geneva, 1211 Geneva, Switzerland

**Keywords:** Pericyte, mechanobiology, confinement, migration, Brownian motion

## Abstract

Pericytes are mural cells of the microvasculature, characterised by their elongated distinct shape. Pericytes span along the axis of the vessels they adhere to, therefore they experience extreme lateral and longitudinal confinement. Pericyte shape is key for their function during vascular regulation and their spatial distribution is established by cell migration during the embryonic stage and maintained through controlled motility in the adult. However, how pericyte morphology is associated with migration and function remains unknown. We use micropatterns to mimic pericyte adhesion to vessels, and to reproduce in vitro the shapes adopted by pericytes in vivo. We show that lateral confinement controls cell shape and produces in vivo-like phenotype. Modelling the pericyte as an incompressible linear elastic material predicts strain and shape of pericytes as a function of lateral confinement. Pericyte kinetics on both laterally confining lanes, and longitudinally constraining motifs is described by dry friction theory. Pericytes are capable of crossing gaps of different sizes. The percentage of crossings is correctly predicted by the likelihood of a fluctuating system to overcome an energy barrier. Our joint experimental and theoretical approach demonstrates the effect of in vivo-like geometrical confinement on pericyte morphology and migration which is accurately described by dry friction theory.

## Introduction

Pericytes are mural cells of the microvasculature, they wrap around microvessels and are in direct contact with endothelial cells (1; 2; 3). Pericytes are characterised by their distinctive shape: a protruding cell body called soma, from which thin and long processes emanate, contains the nucleus and most organelles, while the processes are mainly composed of thin actin filaments (4; 5; 6). Because of the lack of pericyte-specific biomarkers, this unique cell shape, thought to be essential for their function, is used to identify pericytes in vivo (7; 8; 9). Pericyte morphology varies along the microvascular tree (10; 11; 12). On arterioles, pericytes are short ( ∼ 40 *μ*m) and present many encircling processes; on capillaries, in contrast, they are much longer (∼ 150*μ*m) with mainly two longitudinal processes extending on each side of the soma (12). Despite the fact that pericyte morphology plays a pivotal role in their function, the mechanisms that regulate their morphology remain largely unknown, and native phenotypes have not thus far been reproduced in vitro.

Pericyte distribution along the microvasculature follows an archetypal pattern, where they form a distinctively spaced cellular chain, particularly noticeable in the brain (12; 13). To achieve this tightly controlled positioning, pericytes migrate to a specific location during embryonic development (14). This migration is a two-step process, involving process extension followed by nucleus translocation (1). Cells transition to an immobile state subsequently, where they acquire their characteristic morphology. The final distribution ensures a regular coverage of the microvascular tree, where each pericyte covers a specific domain, bound longitudinally by the presence of neighboring pericytes (15; 11). This domain-bound coverage is maintained by a mechanism similar to contact inhibition of locomotion, where pericytes constantly probe their surroundings and retract when touching another pericyte’s process (15; 2). Upon perturbation of the network, for instance after laser ablation of one pericyte, the adjacent pericytes extend their processes drastically to cover the denuded microvessel wall (15). These observations suggest critically regulated cell dynamics with an essential role in cell function, which is likely to depend on cell morphology or their ability to detect boundaries.

Smart in vitro systems, designed to decipher the regulatory pathways underlying basic cell characteristics and functions (such as morphology or migration) allow disentangling the role of the various intricate cues in vivo (16). In vitro models, and more specifically geometrical motifs, have been used to guide cells into the native phenotypes, after the identification of the controlling cue: adhesive lines for muscle cell alignment and differentiation or endothelial cell alignment (17; 18), microstructured surfaces for neuron differentiation and axon elongation (19), and various geometrical motifs to study cell migration (16). The established correlation between the different pericyte morphologies and vessel diameter suggests that geometrical cues such as lateral confinement or curvature, which could both be imposed in in vitro setups, might similarly play a role in directing pericyte phenotype (1; 12).

In addition to custom experimental platforms, theoretical frameworks have been employed to establish physical principles driving cell migration. A class of models have broached the question from a mechanistic point of view, for instance by modelling the protruding force generation by the polymerising actin at the cell front, and balancing it against the friction force generated by focal adhesions at the cell rear (20). Other models have focused on cell-scale dynamics and kinetics. These are often represented by the Ornstein-Uhlenbeck stochastic process (21; 22), which describes the motion of a particle animated by random noise (stemming from biochemical activity such as polymerisation of different proteins) and subjected to friction (stemming from cell adhesion to the substrate). Whereas friction between a moving cell and a solid substrate is often modelled as viscous drag, proportional to the cell velocity, it has been shown that cell motion may be better described by dry, Coulomb-like friction, independent of cell velocity (23). While studies on dimensionally reduced geometrical motifs combined with relevant theoretical frameworks have produced extensive quantitative analyses and revealed deterministic patterns of the shape and motility for many cell types (24; 25; 26; 27; 28), few studies have explored the mechanobiology of pericytes. Many questions such as what drives the specific shape of pericytes, and how to describe their characteristic mode of migration remain unaddressed. For instance, virtually all in vitro studies of pericyte migration have used 2D surfaces, with in plane migration such as the wound healing assay or even out of plane migration with the Transwell assay. In studies conducted in 3D matrices, pericyte migration is poorly controlled and difficult to quantify (29; 30; 31; 32; 33). Here we use micropatterned adhesive motifs to study the effect of lateral and longitudinal confinement on pericyte morphology and motility. We show that lateral confinement, mimicking the diameters of microvessels, controls pericyte shape and produces in vivo-like phenotypes. Modelling the pericyte as an incompressible linear elastic material predicts the strained shape of pericytes as a function of lateral confinement. Pericyte kinetics on both laterally confining lanes and longitudinally constraining motifs is described by dry friction theory. We further demonstrate the ability of pericytes to cross non-adhesive gaps of different sizes, used to model the discontinuous fibronectin patches found in the in vivo matrix. The likelihood of gap crossing is well described by the probability of a fluctuating system to overcome an energy barrier. Overall our results reveal the effect of in vivo-like geometrical confinement on pericyte shape and we present a new theoretical framework that rationalizes pericyte behaviour during migration based on dry friction.

## Results

### Lateral confinement controls pericyte shape

Native pericytes exhibit long thin processes and protruding bodies, while pericytes cultured in vitro on 2D surfaces exhibit polygonal shapes, large surface areas, discoid flat nuclei and prominent actin cables, suggesting a complete phenotype loss (Fig 1Ai) (34; 35; 36). To examine the effect of the confinement imposed by microvessel geometry on pericyte morphology, we cultured human brain vascular pericytes on micropatterned lanes of varying width, coated with fibronectin (Fig S1A). The different widths (5-20 μm) approximate the half-perimeter of native capillaries and arterioles (Fig 1Aii). Pericyte morphology is modified by the confinement: cells present thin long processes, and can be several hundreds of microns long, comparable to in vivo pericyte lengths. Furthermore, a reduction of the number of stress fibers is observed (Fig 1Aiii-iv). Cell morphometrics are quantified using binarized staining images for actin and nuclei. Cell length increases with confinement, from 150 μm for cells on 20 μm lanes to 350 μm for those on 5 μm lanes (Fig 1Bi), scaling as 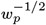 (Fig S1Bi), while the width of pericytes decreases with confinement, following the lane width (Fig 1Bii), scaling as 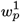 (Fig S1Bii), both replicating in vivo dimensions (12). Cell area also decreases with confinement (Fig 1Biii), scaling as 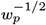 (Fig S1Biii), consistent with the length and width scalings. Finally, the ratio of cell area to pattern area, referred to as coverage, decreases with confinement (Fig 1Biv), as the longer cells exhibiting thin processes found on the narrowest lanes cover a smaller surface. Taken together, these results demonstrate that lateral confinement recapitulates in vivo-like phenotypes in isolated cultured pericytes. (12).

**Figure 1.**
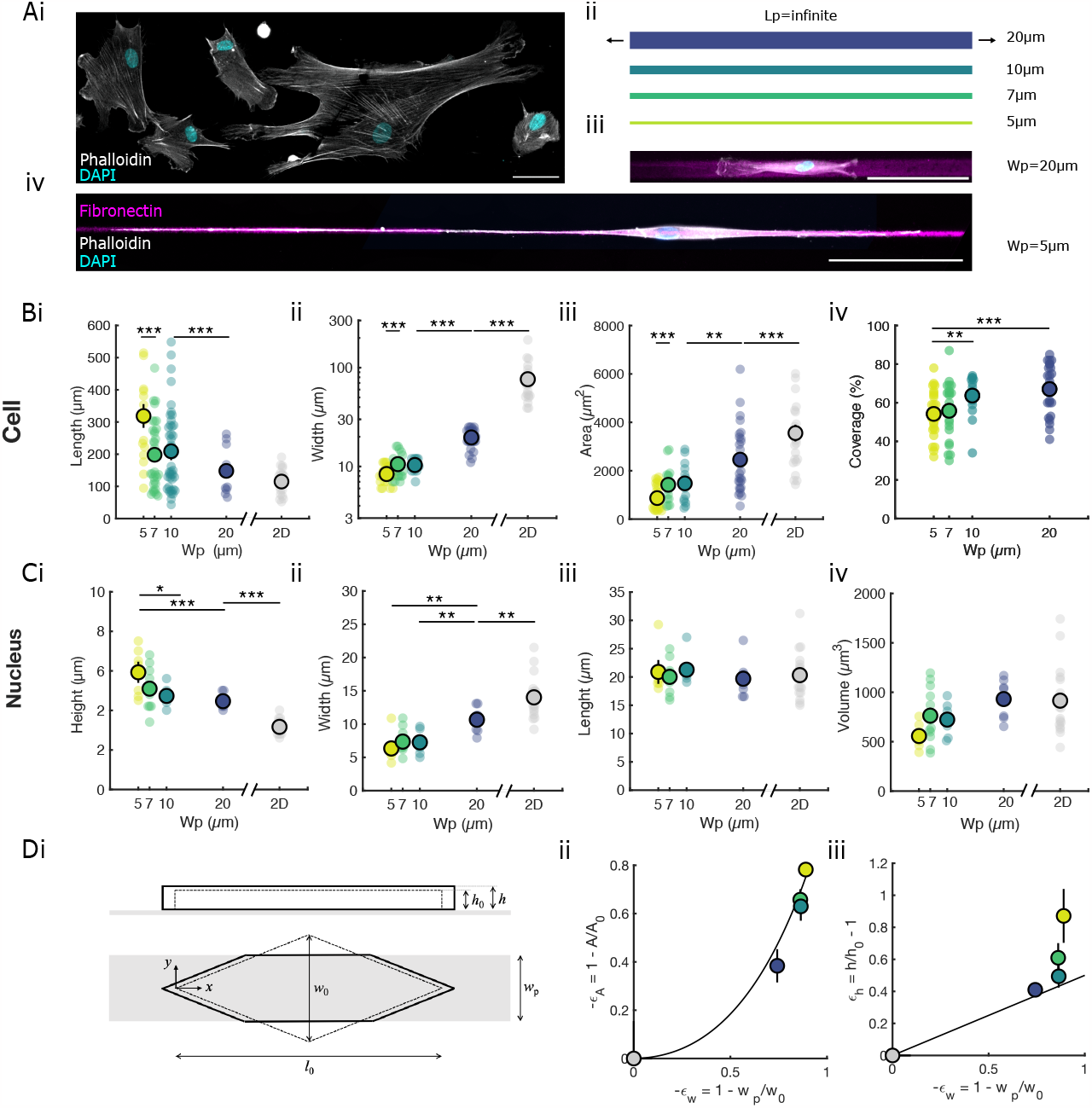
Lateral confinement controls pericyte shape. **(A(i))** 2D culture of pericytes on a fibronectin coated glass surface, immunostained for nuclei (cyan) and actin (white). Scale bar 40 μm. **(A(ii))** Illustration of the pattern lanes with varying width, from 5 μm (light green) to 20 μm (deep blue). **(A)** Pericyte shape on 20 **(iii)** and 5 **(iv)** microns wide lanes, coated with fibronectin (magenta), immunostained for actin (white) and nucleus (cyan). Scale bar 100 μm. **(B)** Mean cell length **(i)**, width **(ii)** and area **(iii)** as a function of pattern width. **(B(iv))** Pattern coverage, defined as the percentage surface of the pattern covered by the cell, as a function of pattern width. **(C)** Height **(i)**, width **(ii)**, length **(iii)** and volume **(iv)** of the nucleus as a function of pattern width. **(D(i))** Idealized shape of a pericyte as a rhomboid (relaxed, dashed lines), deformed by lateral confinement (solid lines). **(D)** Cell area **(ii)** and nucleus height **(iii)** strain as a function of the lateral strain in good agreement with the theoretical prediction (black line). In all panels each dot presents one cell and those with a black contour the mean value.

Another important feature of the native phenotype is the protruding soma, which houses the nucleus. Nucleus height, measured from 3D reconstruction of confocal images, is strongly affected by confinement. While unconfined cell nuclei are discoid and rather flat with a large area (Fig S1C), increasing confinement from 20 to 5 μm doubles nucleus height progressively from 3 to 6 microns (Fig 1Ci). As the nucleus width decreases with the confinement strength (Fig 1Cii), the nucleus height progresses from being smaller than its width to being equally large and tall on the narrowest lanes. This suggests that the nucleus is squeezed laterally by the confinement (Fig S1C). While the length of the nuclei is unchanged across conditions (Fig 1Ciii), the volume of the nuclei trends down with the decreasing widths (Fig 1Civ), suggesting a possible nucleus compression.

We propose a conceptualization of the pericyte as an elastic solid to rationalize the effect of confinement on pericyte shape. In short, we idealize the relaxed shape of the pericyte as a rhomboid of length *l*_0_ and width *w*_0_ in 3D (Fig 1Di, dashed lines). Because the pattern width *w*_p_ is smaller than the relaxed pericyte width *w*_0_, the pericyte is constrained to adapt its maximum width to that of the pattern (Fig 1Di, solid lines) which elongates the pericyte because of the Poisson effect. A simple linear elastic calculation yields a prediction of the pericyte’s areal strain,

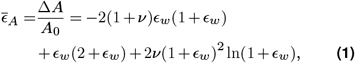

where Δ*A* = *A* − *A*_0_ is the variation of surface area between the confined and relaxed pericyte, *E*_*w*_ = (*w*_p_ − *w*_0_)*/w*_0_ is the confinement strain, and *ν* is the Poisson coefficient. To evaluate Eq. 1, we assume volume incompressibility and take *ν* = 0.5 (37; 38). Based on the assumed rhomboid shape, we estimate the unconfined width as *w*_0_ = 2*A*_0_*/L*_0_. For unconfined pericytes we measure an area *A*_0_ ≈ 3500 *μ*m^2^ and a length *L*_0_ ≈ 115 *μ*m, which yields *w*_0_ ≈ 60 *μ*m. We note that this value *w*_0_ ≈ 60 *μ*m is also the expected confinement threshold, i.e., the maximal pattern width above which we expect pericyte shape to be virtually identical to its unconfined 2D shape. With these parameter values, Fig 1Dii shows a comparison between the predictions of Eq. 1 and the experimental values for the area strain. The good agreement means that the change in dimensions of a laterally confined pericyte is well described by an incompressible, linear elastic solid model. We emphasize that our model has no fitting parameters, as parameters *ν* and *l*_0_ have been determined independently of the data shown in Fig 1Dii.

Our interpretation of the pericyte nucleus being laterally compressed is supported by the linear elastic model predictions. The solid black line in Fig 1Diii shows the expected height variation of the pericyte, *h* − *h*_0_, as a function of the pattern width. The elastic model provides a good description of the experimental observations (dots) for all patterns larger than 7 μm, suggesting that the increase of cell height (which is at least the nucleus height) can be attributed to an elastic Poisson effect in this regime of strain. In contrast, the measured cell height for 5 μm patterns deviates from the prediction of the elastic model, indicating that, under very strong confinement, the overall cellular shape can no longer be fully described by a homogeneous, incompressible, linear elastic deformation of the entire cell body. We attribute this effect to the complex mechanical response of the nucleus (Fig. 1C), which is likely to be both compressible (nuclei volume decreases with confinment) and anisotropic (despite the lateral compression there is no increase in length but a large increase in height).

Overall, the effect of lateral confinement on pericyte morphology can be summarized as follows: it increases length, decreases surface coverage, promotes the soma and process morphology, reduces stress fibers, and increases the height of the nucleus, all of which are markers of the in vivo phenotype.

### Pericyte migration on confined lanes is described by dry friction

One of the distinct behaviors of the pericytes in vivo is their ability to extend their processes over tens of micrometers, and even migrate or “crawl” along the vessel wall (15; 14). We investigate the motility of pericytes on lanes of different widths by following the trajectories of the cell centroid and the extremities of each cell segmented with a Weka-based machine learning algorithm (39). Cells have a variety of motility patterns (Fig 2B), with shorter cells exhibiting a more dynamic behavior, suggesting a correlation between shape, which we showed to be linked to lateral confinement, and velocity.

**Figure 2.**
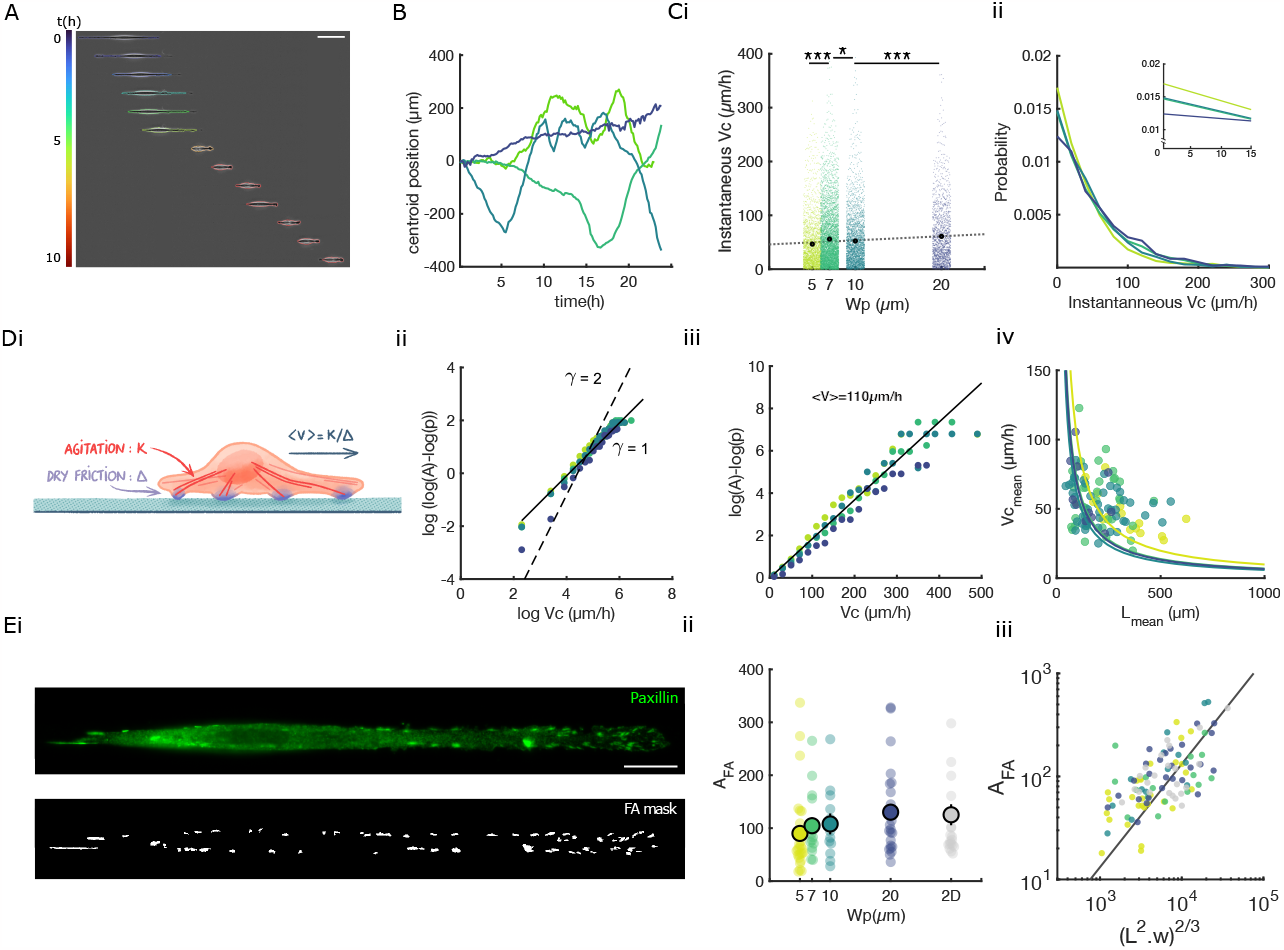
Pericyte migration on confined lanes is described by dry friction. **(A)** Kymograph of a pericyte moving on a 5μm lane over 10 hours with its contour color-coded for time. Scale bar 100 μm **(B)** Cell centroid position for individual cells on four different lane widths, from 5 μm (light green) to 20 μm (dark blue). **(C (i))** Instantaneous velocity of the cell centroid as a function of lane width. **(C(ii))** Probability density of instantaneous cell velocity. The inset zooms on the probabilities for near zero velocities. **(D(i))** Schematic of our proposed conceptualization of pericyte migration, where propulsion stemming from the constant internal agitation is balanced by shape-dependent friction with the substrate. **(D(ii**,**iii))** Probability distribution of the instantaneous velocity of the cell centroid for the four different lane widths, from 5 μm (light green) to 20 μm (deep blue), replotted to highlight the fitted *γ* (i) and the fitted *v*_m_ (ii). The constant *A* corresponds to the experimental probability of having zero velocity, *A* = lim_*v*→0_ *p*(*v*). **(D(iv))** Mean velocity of the cell centroid as a function of cell length, for the four different lane widths, from 5 μm (yellow) to 20 μm (blue). The fitted B coefficients for pattern widths from 5 to 20 are 1.10^4^, 7.10^3^, 6.10^3^ and 7.10^3^μm^2^*/h* respectively. **(E(i))** Immunoflorescence staining of focal adhesions for paxillin and the binarized image. **(E(ii))** Focal adhesion area as a function of lane widths. **(E(iii))** Focal adhesion area as a function of (*L*^2^*w*)^2*/*3^, showing a collapse of the data for the four different lane widths and the 2D substrate, from 5*μ*m (light green) to 20*μ*m (deep blue), with the predicted power law of exponent 1 overlayed (black line). In all panels each dot presents one cell and those with a black contour the mean value.

Indeed, lateral confinement decreases the mean instantaneous cell velocity (Fig 2Ci), which still shows a wide distribution (across two orders of magnitude). The instantaneous cell velocity distribution follows an exponential decay (Fig 2Cii), whose slope depends on the lateral confinement strength: confinement causes a faster decay, in other words, a decreased probability of high velocity and an increased probability of near-zero velocities (Fig 2Cii-inset).

To examine the origin of the probability distribution’s shape and the possible link between cell velocity and cell length, we model pericytes as particles undergoing 1D Brownian motion (Fig 2Di). Each cell is hypothesized to have a fixed level of energy, stemming from internal biological processes such as ATP consumption, akin to the thermal agitation of classical Brownian particles. The internal agitation generates a propulsion force which is then balanced by friction with the substrate (Fig 2Di). This picture can be described by an OrnsteinUhlenbeck statistical process, which is often applied to model cell motion (40; 41; 42). There are two widely used models of friction to describe the energy dissipation due to the relative motion of two surfaces: viscous or dry. In viscous friction, arising for example when a solid moves within a liquid, the force is proportional to the velocity of the object; whereas in dry friction, characteristic of the contact between two solids, the force is independent of velocity. In general, as the two types of friction may be present simultaneously, cell motion can be described by a generalized Langevin equation including both friction terms, as conceptualized by De Gennes (43):

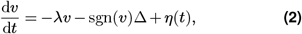

with *λ* the viscous friction coefficient, sgn(*v*) the sign of the velocity, Δ the magnitude of the dry friction force, and *η*(*t*) the Gaussian white noise term. The instantaneous velocity distribution predicted by this model is a Maxwell probability distribution of the form *p* ∼ exp(− *αv*^*γ*^), in line with the exponential shape observed in experiments (Fig 2Cii), where the dominating type of friction will set the value of the exponent: *γ* = 1 for dry friction and *γ* = 2 for viscous friction. We therefore replot the velocity data in a log-log scale and observe that our experimental data is best fitted by *γ* = 1 (Fig 2Dii), demonstrating dry friction between the cell and the substrate. This conclusion is consistent with an earlier report on the existence of exponential tails in the velocity distributions of several cell types, suggesting that cell-substrate friction is better described as dry friction rather than viscous friction (23). From Eq. 2, the following probability distribution of velocities is obtained:

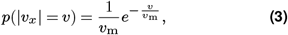

with *v*_m_ the mean velocity, given by (43):

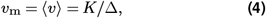

where *K* characterizes the random noise correlation, ⟨*η*(*t*_1_)*η*(*t*_2_)⟩ = *Kδ*(*t*_1_ − *t*_2_), which can be interpreted as a measure of the metabolic agitation of the cell, and Δ is the friction magnitude. Eq. 3 correctly describes the experimentally measured instantaneous velocity distribution with a fitted average velocity of *v*_m_ = 110 *μ*m/h (Fig. 2Diii).

The above description considers the whole cell population together, independent of cell length. To investigate the effect of cell length on velocity, we hypothesize that cells have an intrinsic noise intensity *K*, independent of their shape, but differ in their friction magnitude Δ, which we expect to be proportional to the cell-substrate contact area, Δ = *cLw*, with *c* a constant, *L* the cell length and *w* the cell width, assumed approximately equal to the lane width, *w* ≈ *w*_p_. These hypotheses lead to the following scaling relationship:

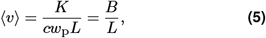

where we define the constant *B* = *K/*(*cw*_p_). Plotting the mean cell velocity ⟨*v*⟩ as a function of the mean cell length *L* reveals a hyperbolic function, matching the prediction of Eq. 5 (Fig. 2Div). Shorter cells are subjected to smaller friction, leading to larger velocities. Overall, as shown in Fig. 2D, Eqs. 3 and 5 provide a good representation of our experimental data with fitted parameter values *v*_m_ ≈ 110 *μ*m/h, corresponding to the observed average instantaneous pericyte velocity, and *B* ≈ 1.1 *·* 10^4^ *μ*m^2^/h. We can rationalize the latter value by noting that, according to our mathematical description, *B* scales as the product of the average cell velocity and length, *B* ∼ ⟨*v*⟩ *⟨L⟩* ≈ 100 *μ*m*/*h *·* 100 *μ*m = 10^4^ *μ*m^2^*/*h.

To probe our hypothesis that friction scales with the cell area, we investigate the dependence between pericyte velocity and pattern width. We find the velocity-length relationship to be independent of pattern width (Fig 2Div), in disagreement with our initial friction scaling assumption, which would require the relationship velocity-length to vary with pattern width (Eq. 5). We thus investigate an alternative hypothesis where friction would scale with focal adhesion area, rather than total cell area.

To this end, we image focal adhesions (FAs), the principal anchoring sites between cells and their substrate (Fig 2Ei). The FAs are clearly visible and are localized mostly at the cell periphery. The total FA area *A*_FA_ is independent of the width of the pattern (Fig 2Eii), as evident by the near zero exponent for the scaling with pattern width (Fig S2A), due to the fact that pattern width affects both cell width and length in opposite ways. The conserved quantity for the focal adhesion area is neither the surface nor line density as both reveal a nonnull scaling with pattern width (Fig S2Bi,ii). If we assume that the total FA area scales with both the length and the width of the cell, we derive that *A*_FA_ ∼ *L*^*α*^*w*^*β*^. Using our previously established scaling for the length 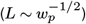 and the width 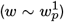, we get 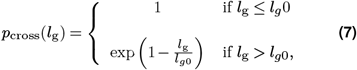, giving us *α* = 2*β*. Dimensional analysis of *A*_FA_ ∼ *L*^*α*^*w*^*β*^ also gives us that *α* + *β* = 2. We then derive that *A*_FA_ ∼ (*L*^2^*w*)^2*/*3^. Plotting the total FA area as a function of (*L*^2^*w*)^2*/*3^ reveals the expected linear relationship and a collapse of the data from the various widths (Fig 2Eiii). This scaling is consistent with the FA being mostly at the periphery, with the conserved quantity being close to a line density, with a small contribution of the cell width. Therefore, to correctly describe the experimental data, Eq. 5 should be rewritten as:

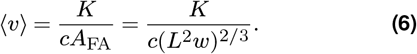

### Constrained length caps cell velocity

In physiological conditions pericytes are seldom free to roam on infinite lanes; there is evidence for their territorial and cell contact avoidance behavior which constrain them to remain in delimited zones (15; 2). Exploiting our highly controllable in vitro system, we investigate whether an additional longitudinal constraint would alter pericyte dynamics. We fabricate patterns with the constant width of 10 *μ*m but with finite lengths from 50 to 350 *μ*m, similar to the territory spanned by a pericyte in vivo (Fig 3A). A longitudinal constraint alters pericyte shape: on longer patterns a higher fraction of cells present in vivo-like morphologies, namely long processes, while shorter patterns decrease the length of processes and increase the number of cells spreading over the total surface of the pattern. Cells present a variety of trajectories (Fig 3Bii), depending on their length, consistent with observations on continuous lanes: shorter cells are quicker and longer cells are slower. But when the length of the cells is near the length of the pattern, cells are immobilized by the lack of available free surface and their velocity drops towards zero. The shorter the pattern, the higher the fraction of cells spreading over the whole pattern length, leading to an increased probability of near-zero velocities (Fig 3Ci, inset). Nevertheless, the probability distribution of the instantaneous velocity follows a similar decaying exponential (Fig 3Ci) as the one found on infinite lanes, suggesting that underlying cell dynamics are unaffected.

**Figure 3.**
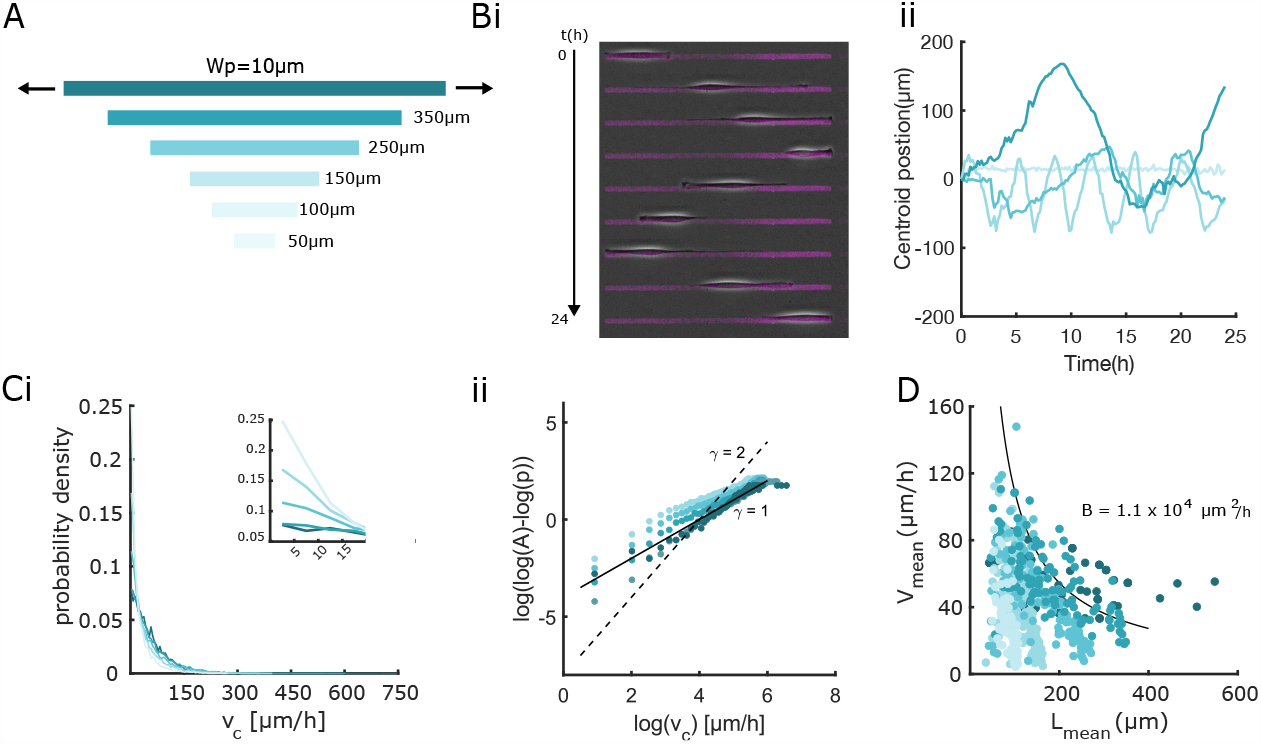
Constrained length caps cell velocity. **(A)** Illustration of the patterns of different lengths over a range of 50 μm to 350 μm. **(B(i))** Kymograph of a cell migrating on a pattern of length 350 μm over 24 hours. **(B(ii))** Examples of cell centroid trajectories, one for each pattern length, from 50 μm (light blue) to 250 μm (dark blue). **(C(i))** Probability distribution of the instantaneous velocity of the cell centroid, for different pattern lengths from 100 μm (light blue) to infinite lanes (dark blue), inset showing the different probabilities of near-zero velocities. **(C(ii))** Probability distribution of the instantaneous velocity of the cell centroid, for the different pattern lengths, from 100 μm (light blue) to infinite lanes (dark blue), replotted to highlight the fitted *γ*. The constant *A* corresponds to the experimental probability of having zero velocity, *A* = lim_*v*→0_ *p*(*v*). **(D)** Mean velocity of the cell centroid as a function of cell length, for the different pattern lengths, from 100 μm (light blue) to infinite lanes (dark blue), with the theoretical hyperbolic curve fitted to the infinite lanes (black line).

Indeed, the distribution is still best fitted by *γ* = 1, despite a slight shift observed for strongly constrained cells, confirming that the constrained cell motion is driven by dry friction theory (Fig 3Cii). Plotting the mean cell velocity as a function of mean cell length confirms that for each pattern length, cells follow the same hyperbolic decrease as on infinite lanes, with a similar *B* coefficient, until the cell length nears the pattern length, at which point their velocity drops toward zero (Fig 3D, S3).

### Gap crossing and constraint release

Another feature of pericyte migration in vivo is their ability to overcome discontinuities in adhesion or even to bridge two separate capillaries (44; 2). To verify whether cells in our in vitro model could replicate this behavior we produced patterns of finite length (from 50 *μ*m to 350 *μ*m) and of a constant width of 10 *μ*m, but separated by gaps of different sizes; 5, 10, 15, 20 and 30 *μ*m (Fig 4A). Cells are found to successfully bridge the gaps, either by spreading to the neighboring pattern, with part of the cell remaining on the initial pattern (Fig 4A), or by crossing, with the whole cell moving to the adjacent pattern (Fig S4). The probability of a pericyte bridging a gap decreases with increasing gap size until it reaches zero for gaps of 30 *μ*m (Fig 4Bi).

**Figure 4.**
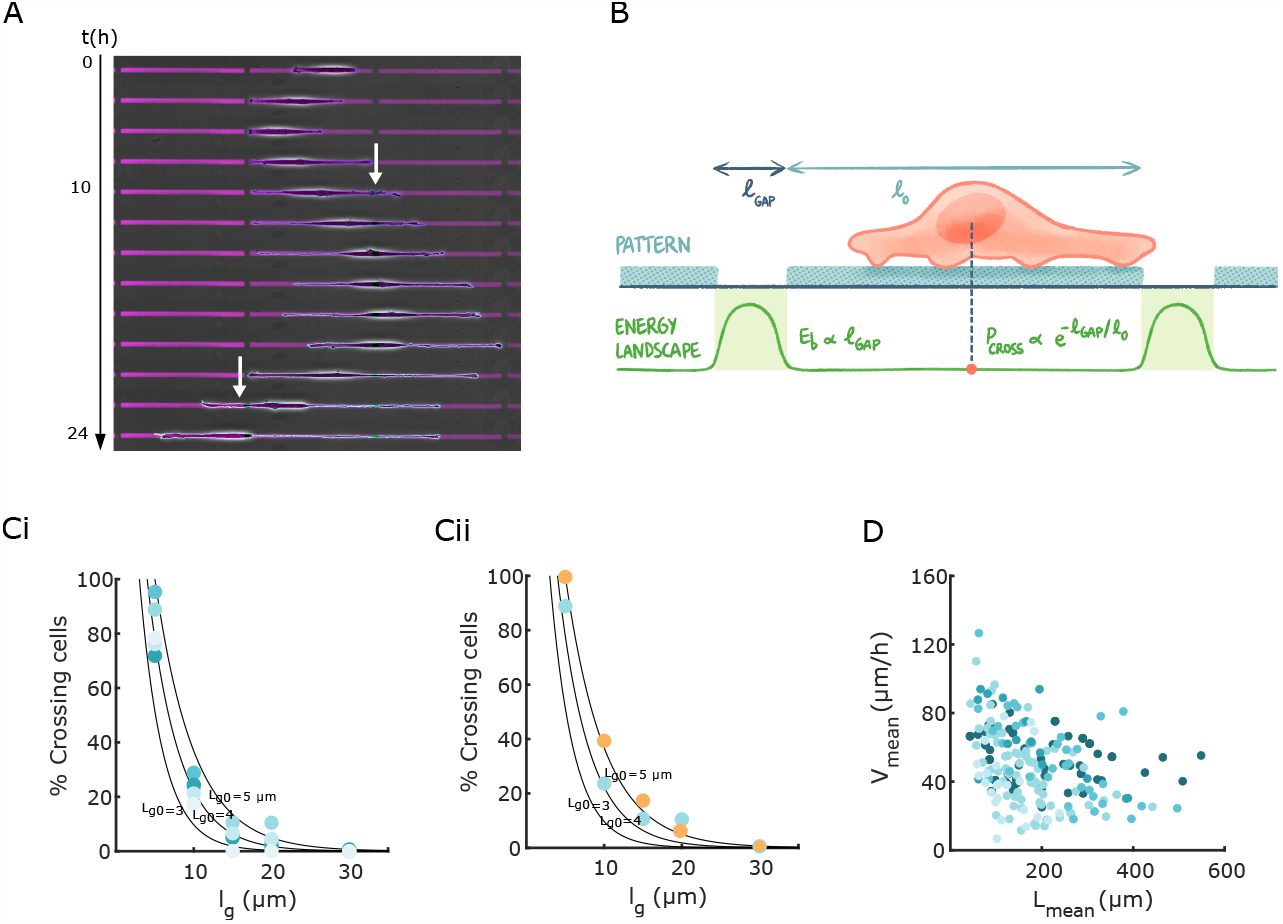
Gap crossing and constraint release. (A) Kymograph of a pericyte migrating on patterns of length 250 μm over 24 hours, showing two subsequent bridgings of a gap of 10 μm (white arrow), with the final cell spreading over three patterns. **(B)** Schematic of our proposed conceptualization of gap crossing by pericytes, where the non-adhesive gaps represent an unfavorable surface for migration modeled as an energy barrier, whose value depends on the gap length. **(C(i))** Percentage of crossing cells as a function of gap size, for different pattern length from 100 μm (light blue) to 350 μm (dark blue). The curves are fits of the theoretical probability distribution (Eq. 7). **(C(ii))** Percentage of crossing cells as a function of gap size for two different pattern widths, 10 μm (blue) and 20 μm (orange). **(D)** Mean velocity of the centroid of cells which have successfully crossed the gaps, matching that of cells on infinite lanes, for different pattern length from 100 μm (light blue) to infinite lanes (dark blue), independent of gap size.

We conceptualize these observations by our description of pericytes as a statistical system and by assimilating the nonadhesive gap to an energetic barrier (Fig 4B). The likelihood of a fluctuating system to overcome an energetic barrier decreases exponentially with the height of the barrier, which suggests the following form for the probability distribution of crossing a gap:

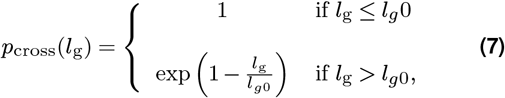

where *l*_g_ is the length of the gap and *l*_*g*_ 0 corresponds to the longest gap that cells virtually always traverse, which can be viewed as a fitting parameter. Fig 4C shows that Eq. 7 is in good agreement with the experiments for a choice of *l*_*g*0_ between 4 and 5 *μ*m, a value consistent with our interpretation of this parameter as the longest gap that cells should always cross. We find no significant effect of the pattern width on the percentage of cells crossing the gaps (Fig 4Cii), consistent with our previous result of pericyte dynamics being insensitive to pattern width (Fig. 2D), due to the conserved total area of focal adhesions (Fig. 2Eii).

Interestingly, plotting the velocity of crossing cells as a function of the cell length shows the same hyperbolic dependency as on the infinite lines (Fig 4D), rescuing the decaying tails found in the case of constrained cells by offering free surface to migrate on, more specifically the neighboring pattern. This further suggests that the cell dynamics is not significantly affected by the short non-adhesive regions.

## Discussion

In this study we use a highly controllable 1D in vitro system combined with a theoretical framework to reveal the impact of geometrical confinement on pericyte shape and dynamics. We demonstrate that lateral confinement induces in vivo-like morphology in pericytes, where both the length of pericytes and the height of the soma increase as the lanes become narrower. These changes are similar to observations in studies conducted in vivo, where thin-strand pericytes on capillaries are much longer than cells found on larger microvessels (1; 15). Another important feature, which also allows to differentiate pericytes by their morphology, is their coverage of the vessel walls. Cells on wider lanes cover a larger surface of the patterns, reproducing the various vessel coverage by pericyte subtypes in microvasculature: thin-strand pericytes found on capillaries cover a smaller surface of the microvessel compared to ensheathing or stellate pericytes found respectively on pre-capillary arterioles and post-capillary venules (1; 15). We show that modelling the cell as an elastic isotropic incompressible material predicts the cell behavior quite accurately. The theoretical curve predicting the dependency of areal strain on the lateral strain fits well our experimental data. This model presents a first approach to understand pericyte mechanics, but further models could be explored to increase the complexity and reach the in vivo situation. However, it informs us that shape adaptation in pericyte is impacted by an underlying elasticity, probably arising from its cytoskeleton. Moreover, the observed incompressibility indicates that pericyte volume is conserved through shape adaptations. The change in cell height, based on the same assumptions, follows the model prediction for small deformations before diverging for increasing confinements.

The substrate pericytes adhere to in vivo is not a homogeneous layer of one type of protein, but rather a complex matrix of multiple proteins, where laminin is the major component forming a discontinuous layer, punctuated with patches of fibronectin (45; 46; 44). In an in vitro study using micropatterned lanes and dots, pericytes were shown to have a marked preference for fibronectin over laminin (47). Although the authors did not study cell shape, their results suggest that matrix proteins can affect cell adhesion by changing focal adhesions and are therefore likely to tune cell shape and migration dynamics (48). Varying cell adhesion through the type and spatial distribution of adhesion proteins, in addition to geometrical confinement, could provide further control over pericyte phenotype within in vitro settings.

While a number of studies have used 1D patterns as a relevant tool to investigate cell migration in highly controllable conditions, only few studies have examined pericyte migration in vitro. They have either focused on single cells on 2D surfaces, where the specific phenotype of pericytes is lost, or on the collective migration of pericytes in a wound healing assay, where pericytes are in a monolayer which is far from in vivo conditions (29; 30). As for 3D assays, one has used a transwell setup, which does not allow to access the dynamics or shape, and another is a vessel-on-chip or vessels embedded in a 3D hydrogel where conditions are too complex to dissociate cues (31; 32; 33). We developed a novel configuration that mimics the 1D migration of pericytes found in vivo, by allowing them to migrate along infinite or constrained lanes, which revealed an anticorrelation between cell length and speed of migration.

We have demonstrated that the classical Ornstein-Uhlenbeck statistical framework, widely used to describe cell motility, can be adapted to characterize pericyte motility on laterally confining micropatterns. Application of this statistical physics framework to pericytes requires certain modifications to capture specific features of pericyte migration. First, unlike the common description of cell-substrate interaction as viscous friction, proportional to cell velocity, our experimental data suggest the use of a dry friction term, independent of velocity. Second, friction is proportional not to the total pericyte area but rather to the area of focal adhesions, suggesting that the dry friction effect arises from focal adhesion dynamics. Interestingly, the observed dynamics appear to rely on robust cellular processes that conserve the cell velocity distribution through a wide array of confinement geometries, including continuous and discontinuous micropatterns of different lengths and widths. A universally observed feature in our experiments, shared with many other cell types, is the inverse correlation between cell length and width (26). This feature is explained by our model by considering that all cells share an intrinsic level of activity, independent of their shape, whereas cell-substrate friction depends on cell shape. Thus, pericytes can adopt a shorter and highly migratory phenotype, which allows them to explore the environment, and then randomly switch to a longer and sessile phenotype to cover a defined portion of a microvessel. This is reminiscent of the transition from short migratory cells, found in the embryo during microvasculature formation, to long quiescent and immobile cells, found in the adult microvasculature, which can revert to a more motile phenotype under certain stresses (1). What regulates this transition and the role mechanical and environmental stimuli could play in it remains to be elucidated.

## Methods

### Micropatterning substrates

Motifs are fabricated using Primo, a maskless lithography micropatterning system (Alvéole) mounted on an inverted microscope (Nikon Eclipse Ti2). Glass-bottomed FluoroDishes (WFD3523) are plasma-activated for 45 seconds (Harrick plasma), prior to surface passivation with a blocking solution of Pll-g-PEG (poly-L-lysine gpolyethylene glycol), (Susos) at 0.1 mg/mL for one hour at room temperature. The substrate is then washed three times with ultrapure water (Milli-Q) while avoiding surface drying. The substrate is covered with photoinitiator (PLLP) before being transferred to the microscope stage. Designed motifs are projected through the Primo Digital Micromirror Device (DMD) using Leonardo software (Alvéole). A precalibrated dose of 1200mJ/mm2 of 375nm UV light pathing through a 20x objective (Nikon, Plan fluor, NA = 0.45), activates the photoinitiator which degrades the Pll-gPEG at the desired locations. After patterning, the photoinitiator is washed off with ultrapure water, the surface is coated with fibronectin at 50μg/ml (Sigma Aldrich) mixed with a small quantity of fluorescent fibrinogen (647, Invitrogen) for 15 min at room temperature. The surface is once again washed with ultrapure water. The quality and the size of produced patterns is checked using fluorescence microscopy. Patterned substrates could be used either immediately after preparation or up to 24 hours after when stored at 4°C.

### Cell Culture

Human Brain Microvascular Pericytes (Sciencell), in passages 2-6 are cultured in pericyte growth medium (Sciencell) at 37°C in a humidified incubator at 95% air and 5% CO2. To promote cell adhesion, culture flasks are coated with PolyLLysine (0.01%, Sigma Aldrich) before each passage. Upon 80% confluence, cells are detached using trypsin (Gibco, Life Technologies) resuspended at 6000 cells/ml and then directly seeded on either the patterned surfaces or 2D control surfaces. Sample are incubated for one hour at 37°C, the nonadherent cells are washed out and fresh medium is added, before the sample is returned to the microscope stage.

### Time-lapse Imaging

Live recording of cell motion is performed with an automated inverted microscope (Nikon Eclipse Ti2) controlled by the NIS software (Nikon), equipped with temperature and CO2 regulation. Images are acquired with a 10x objective (Nikon Plan Fluor NA = 0.30) for 24h at 10 min intervals. Multiple fields of view are chosen to follow cells during the 24h.

### Immunostaining

Samples are fixed with 4% paraformaldehyde (Thermo Fisher) in PBS for 15 min. After 15 minutes of permeabilization in a solution of 0.25% Triton, the sample is covered with 3% bovine serum albumin (BSA) in PBS solution for one hour at room temperature for blocking. After the washing steps, the sample is incubated with the primary antibody, mouse anti-paxillin (1:200, MA5-13356, Thermo Fisher) for one hour at room temperature. To detect focal adhesions the sample is incubated with Alexa Fluor 488-conjugated donkey anti-mouse antibody at 1/400 (ab150105, Abcam) for an hour. Nuclei and actin staining are performed with DAPI and phalloidin at 1/800 and 1/400 in PBS respectively. Immunostained sample are imaged with an epifluorescence inverted microscope (Nikon Eclipse Ti2) (with a Crest X-Light confocal system for nuclei imaging) with a 40X water-immersion objective (N40XLWDNIR, Nikon).

### Image processing

To produce a precise detection of cell shape for moving cells, a custom-made Fiji macro is used. Briefly, a classifier is trained with Weka Segmentation plugin allowing to choose different selected features on a set of images (39). Then a binary image is produced by thresholding and cell shape metrics are extracted from implemented particle analysis. The binary mask of the cell shape is fed to a custom-written script on Matlab to detect the centroid position and the two extremities of the cell to compute cell length and cell velocity. In the case of immunostaining images, thresholding and fitting shapes such as rectangles or ellipses to the objects by hand are used to extract cell shape metrics. All data is processed and plotted using MATLAB.

### Focal adhesion analysis

FA morphometric analysis is performed using a custom-made Fiji macro (49). Briefly, a blurred image of the FAs is subtracted from the original images. Thresholding is used to obtain binary images, eroding is applied to help minimize pixel noise. Morphometric descriptors are extracted using particle analysis.

### Statistical Analysis

All analyses are based on at least three independent experiments, each condition contains at least 20 cells. All data are plotted with the standard error of the mean (SEM). An unpaired Student t-test is used for significance testing between two conditions. Statistical tests were performed using Matlab. *** denotes p < 0.001, ** denotes p < 0.01, and * denotes p <0.05.

## Acknowledgments

We are grateful to Dr Mathieu Dedenon for insightful discussion and to Dr Naïs Coq for illustrations. This work was supported by a doctoral fellowship from Ecole polytechique to A.S., the Human Frontier Science Program (HFSP) RGP0032/2022 to P.J.S., by the EMBO fellowship ALTF 886-2022 to C.A.D. and by Région Ile de France through Dim Elicit program, Plantuidics grant to A.B.

## Supplementary Information

**Figure S1.**
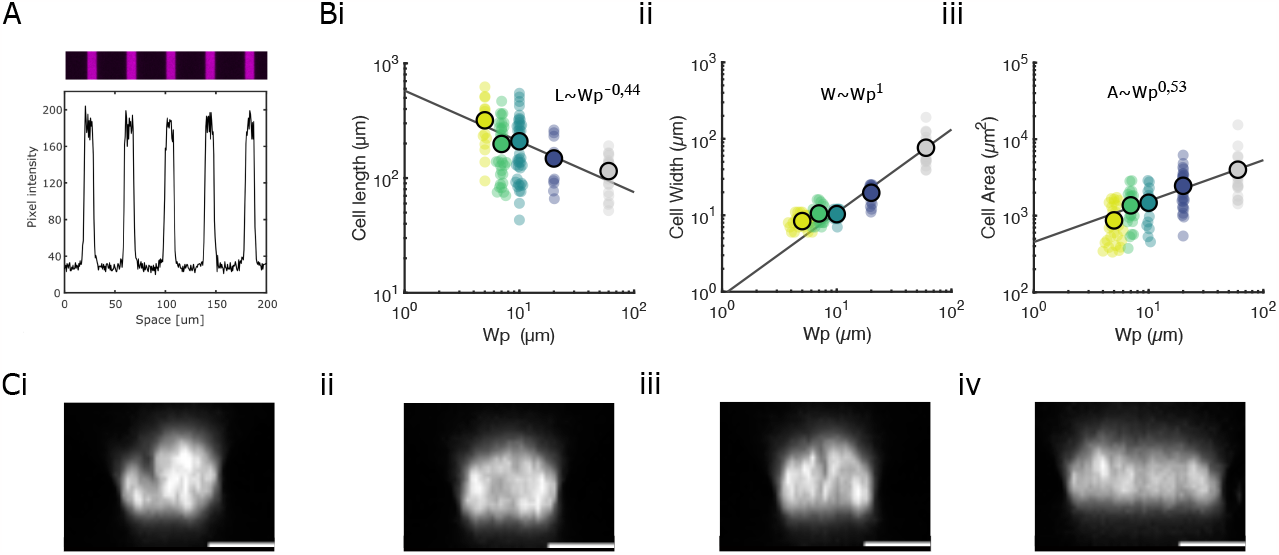
Lateral confinement controls pericyte shape. **(A)**Fluorescent image of fibronectin lines of 10 μm width (top) with its intensity profile (bottom) denoting pattern sharpness. **(B)** Cell length **(i)**, width **(ii)** and area **(iii)** as function of pattern width, fitted with a power law (black line). **(C)** Representative cross-sectional images of pericyte nuclei, stained with DAPI, for pattern widths of 5 **(i)**, 7 **(ii)**, 10 **(iii)** and 20 **(iv)** μm. Scale bar 5 μm.

**Figure S2.**
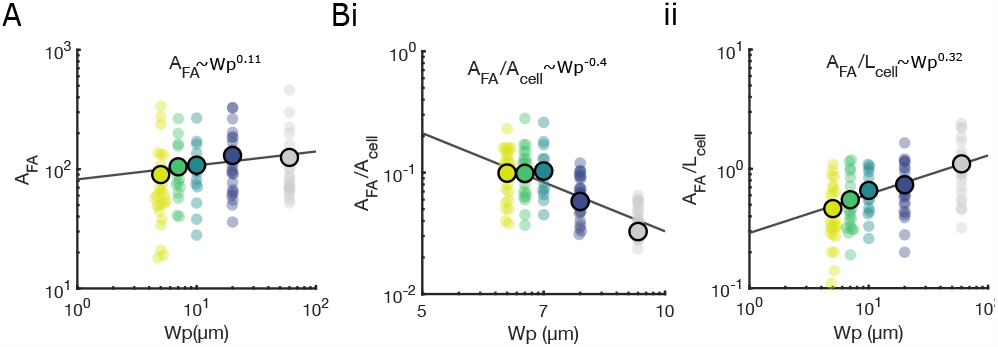
Pericyte migration on confined lanes is described by dry friction. **(A)** Focal adhesion area as function of pattern width, fitted with a power law (black line), revealing a near zero exponent. **(A)** Focal adhesion area normalized by the cell area **(i)** and the cell length **(ii**) as function of pattern width, fitted with a power law (black line), revealing non-null exponents.

**Figure S3.**
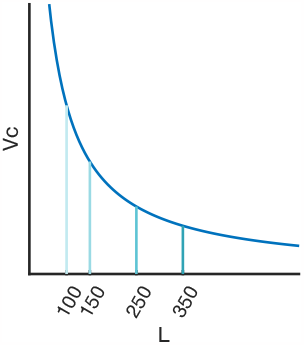
Constrained length caps cell velocity. Schematic of the velocity of the cell centroid as a function of cell length, showing the hyperbolic relationship found for infinite lanes (dark blue), followed by the cells on the different pattern lengths, until the cell length reaches the pattern length and their velocity drops to zero.

**Figure S4.**
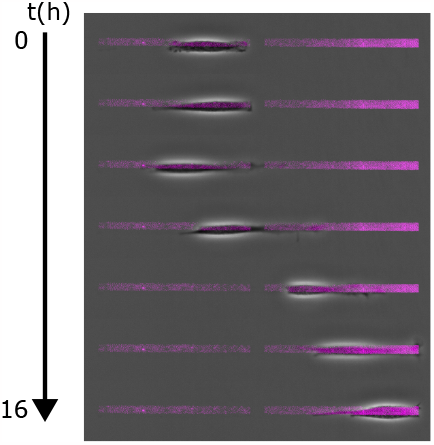
Gap crossing and constraint release. Kymograph of a pericyte migrating on patterns of length 250 μm over 16 hours, showing a crossing of a gap of 10 μm, with the cell having left the initial pattern entirely.

